# Pyramidal neuron synapses in M2 exhibit properties intermediate between prefrontal cortex and M1 synapses

**DOI:** 10.64898/2026.07.06.736741

**Authors:** Abdulmajeed Yarim, Simone Brachtendorf, Hartmut Schmidt, Grit Bornschein

## Abstract

Motor planning and control is executed by different motor areas within the neocortex. Despite their distinct functions these areas are built by the same archetypes of neurons as the rest of the cortex, with the pyramidal neurons (PNs) as their principal building blocks. Recent results suggest that the synapses of the PNs are modeled and adapted to their required functions in an area specific manner. PN synapses in a cortical area engaged in higher order functions, the prefrontal cortex (PFC), were found to operate with loose microdomain calcium-influx-to-release coupling and showed short-term facilitation, whereas synapses processing sensory information in a lower order cortical area, the primary somatosensory cortex (S1), featured tight nanodomain coupling and showed short-term depression. In the present study, we asked for the functional coupling configuration of an intermediate processing area. We focused on PN synapses in the premotor cortex M2 and compared their properties to those of PN synapses in the primary motor cortex M1. In both areas we found tight nanodomain coupling and high release probability, but a significant difference in short-term plasticity. Synapses in M1 showed paired-pulse depression similar to S1. In contrast, synapses in M2 exhibited paired-pulse facilitation. Our data suggest that this facilitation results from an accelerated recruitment of synaptic vesicles to the readily releasable pool from an enlarged replenishment pool. Thus, PN synapses in M2 appear to have properties intermediate between those in PFC and M1.

**Significance Statement:** Neocortical areas perform diverse computations despite being composed of the same principal neuronal cell types. How synaptic properties are adapted to these functional demands remains poorly understood. We show that L2/3–L5 pyramidal neuron synapses in the secondary motor cortex (M2) combine features of the primary motor cortex (M1), including tight nanodomain Ca²⁺-influx-to-release coupling and high release probability, with paired-pulse facilitation, previously associated with higher-order cortical areas such as the prefrontal cortex (PFC). Our data suggest that this facilitation is mediated by an enlarged replenishment pool and accelerated vesicle recruitment rather than differences in coupling architecture, highlighting vesicle pool organization as an additional mechanism of area-specific synaptic specialization. Thus, L2/3–L5 pyramidal neuron synapses in M2 exhibit intermediate properties between M1 and PFC.

## Introduction

Synapses are highly specialized signaling units that are adapted to the computational demands of the neuronal circuits in which they operate. Consequently, synapses differ markedly in their efficacy, reliability, and capacity for short-term plasticity (STP). During the first release process presynaptic efficacy depends on the size of the readily releasable pool (RRP) of vesicles, the Ca^2+^ influx (Mintz et al., 1995; Borst and Sakmann, 1999; Thanawala and Regehr, 2013), the Ca^2+^ sensitivity of the release machinery (Schneggenburger and Neher, 2000; Sakaba, 2008; Bornschein and Schmidt, 2018; Bornschein et al., 2025) and the coupling distance (CD) between voltage-gated Ca^2+^ channels (VGCCs) and the release sensor (Bucurenciu et al., 2008; Eggermann et al., 2011; Scimemi and Diamond, 2012; Vyleta and Jonas, 2014; Nakamura et al., 2015; Bornschein et al., 2019; Schwarze et al., 2026). Together, these factors determine the vesicular release probability (*p*_N_). During repetitive activity, synaptic strength is additionally shaped by the vesicle pool organization and the kinetics of vesicle recruitment (Doussau et al., 2017; Vandael et al., 2020).

Mature synapses specialized for the transmission and processing of high-frequency coded sensory information typically operate with nanodomain coupling (Fedchyshyn and Wang, 2005; Schmidt et al., 2013; Baur et al., 2015; Nakamura et al., 2015; Bornschein et al., 2019), whereas highly plastic synapses operate with microdomain coupling (Vyleta and Jonas, 2014). Although neocortical areas are composed of a common repertoire of neuronal cell types, recent studies suggest that the presynaptic nanoarchitecture of pyramidal neuron (PN) synapses is adapted to the functional requirements of individual cortical areas. In the primary somatosensory cortex (S1), PN synapses specialized for rapid and reliable sensory processing operate with tight nanodomain coupling between a few VGCCs and the Ca^2+^ sensor (Bornschein et al., 2019; Bornschein et al., 2020). In contrast, analogous PN synapses in a higher-order association cortex, the prefrontal cortex (PFC), exhibit looser coupling involving multiple VGCCs (Schwarze et al., 2026).

These findings suggest that synaptic CDs may be systematically related to the computational role of a cortical area. However, it remains unknown how synapses are organized in secondary cortical areas that occupy an intermediate position between primary and higher-order association cortices. Furthermore, it is unclear whether tight nanodomain coupling is a specialization of sensory pathways or represents a more general principle of cortical information processing.

To address these questions, we focused on the cortical motor pathway. Voluntary movements are initiated in higher motor centers involved in the planning and adaptation of motor behavior. In contrast to sensory systems, where information flows from primary sensory to higher-order cortical areas, motor information is transformed in the opposite direction, from higher-order frontal areas toward the primary motor cortex (M1) (Kandel, 2000). Within this hierarchy, the secondary motor cortex (M2) receives input from the PFC and contributes to motor planning and action selection before signals are progressively transformed into execution-related motor commands (Barthas and Kwan, 2017; Yang and Kwan, 2021).

We hypothesized that the functional specialization of M2 and M1 is already reflected at the level of individual synapses. Specifically, we asked whether the premotor cortex M2 exhibits loose microdomain coupling and short-term facilitation similar to the PFC and whether M1 synapses are tightly coupled and depressing similar to synapses in S1. We addressed this issue at synapses between PNs in layer 2/3 (L2/3) and layer 5 (L5) in M2 and M1 using patch-clamp recordings, EGTA-AM experiments, multiple-probability fluctuation analysis and high-frequency train stimulations. Surprisingly, we found tight calcium-influx-to-release coupling and high *p*_N_ in M2 as well as in M1. However, L2/3-L5PN synapses in M2 showed paired-pulse facilitation as opposed to paired-pulse depression in M1. This facilitation was associated with faster vesicle recruitment to the RRP and a larger replenishment pool (RP). Thus, L2/3-L5PN synapses in the premotor cortex M2 exhibit intermediate properties between those of the higher-order association cortex PFC and the primary motor cortex M1.

## Results

### Differential short-term plasticity between PN synapses in M2 and M1

To investigate if the functional properties of synapses differ within the motor pathway we focused on glutamatergic synapses of neurons in L2/3 projecting onto L5PNs in premotor cortex M2 and in primary motor cortex M1.

EPSCs were recorded from whole-cell patch-clamped L5PNs and evoked by minimal extracellular stimulation in L2/3 (**Figure 1A**). The stimulation pipette was placed close to the PN somata in upper layer L2/3, and the stimulation intensity was kept low, making it likely that only the connections between L2/3PNs and L5PNs were stimulated (see Methods for details). In response to paired stimuli at a frequency of 50 Hz we found robust paired-pulse facilitation (PPF) in M2 with a median paired-pulse ratio (PPR) of 1.4 (IQR: 1.15-1.61, n=73). In M1 on the other hand, short-term plasticity (STP) was characterized by paired-pulse depression (PPD) with a median PPR of 0.9 (0.80-1.03, n=15; **Figure 1B-D**).

**Figure 1.**
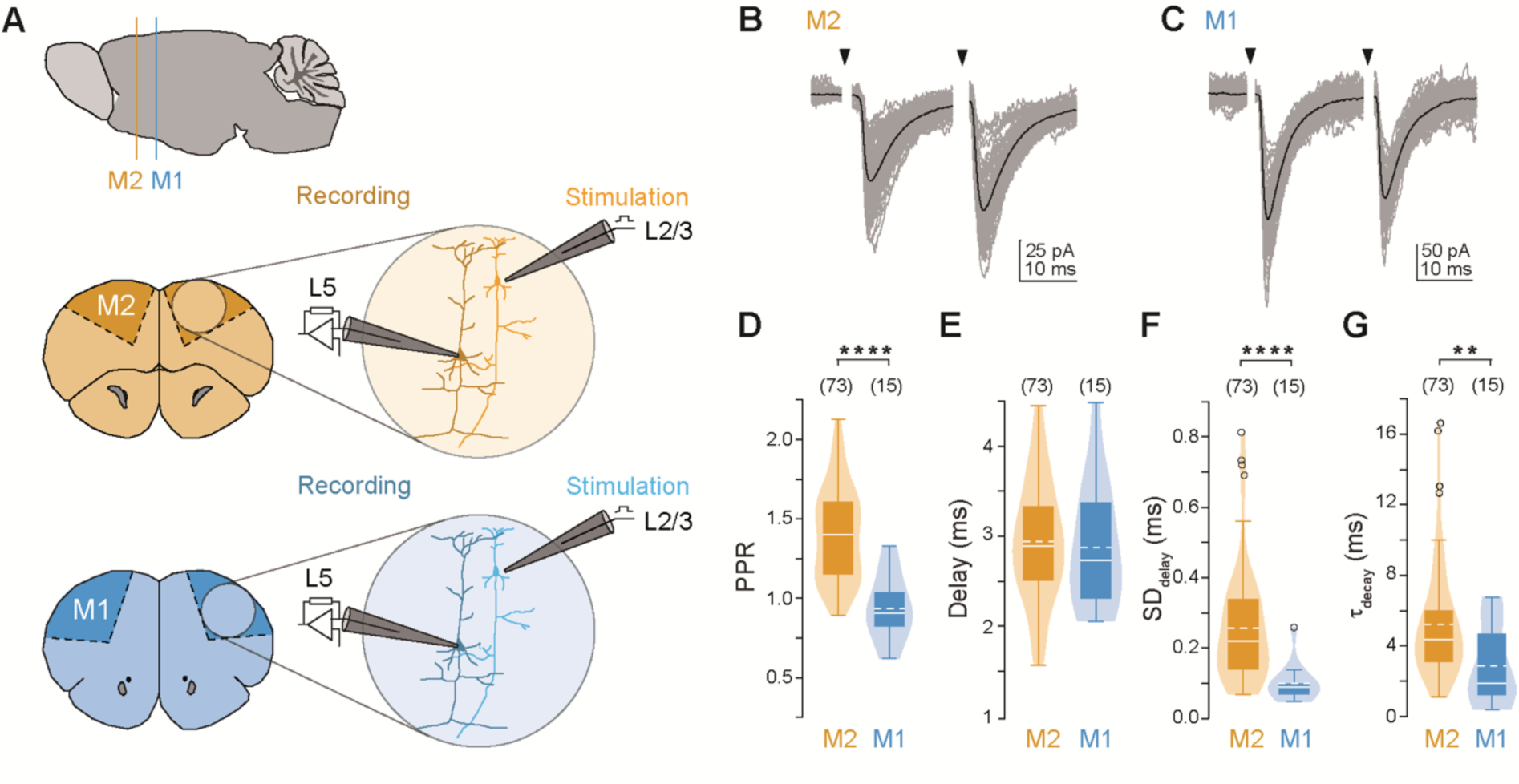
Comparison of short-term plasticity between M2 and M1 cortical regions. (**A**) Schematic representation of the mature mouse brain (*top*) and the positions at which coronal sections were made from premotor cortex (M2, orange; *middle*) or primary motor cortex (M1, blue; *bottom*). Electrophysiological whole-cell recordings were performed in acute slices from layer 5 pyramidal neurons (L5PNs) after extracellular stimulation of presynaptic neurons in layer 2/3 (L2/3). (**B, C**) Representative whole-cell recordings of EPSCs in L5PNs in M2 (**B**) and M1 (**C**). EPSC amplitudes (individual recordings in gray, average in black) were evoked by extracellular stimulation in L2/3 with paired stimuli with an interstimulus interval (ISI) of 20 ms, indicated by the black arrowheads (stimulation artefacts are blanked for clarity). (**D-G**) Quantification of paired-pulse ratio (PPR, **D**), delay (**E**), standard deviation of the delay (SD_delay_, **F**), and decay time constant (τ_decay_, **G**) for M2 (orange) and M1 (blue): Violin plots show the distribution of each parameter, with box plots inside indicating the median (solid line), interquartile range (IQR, box), mean (dashed line), outliers (shown as dots) and number of experiments (in brackets). M2 exhibited significantly higher PPR (****P<0.0001, MWU), SD_delay_ (****P<0.0001) and τ_decay_ (**P=0.003) compared to M1, while the synaptic delay (P=0.556) was similar to M1.

In order to test whether postsynaptic effects may lead to an underestimation of the PPR, we applied the competitive, low-affinity α-amino-3-hydroxy-5-methyl-4-isoxazolepropionic acid receptor (AMPAR) antagonist kynurenic acid (Kyn) to relief postsynaptic saturation and desensitization. In the presence of Kyn, PPRs were not significantly different from the above values neither in M2 (1.19, 1.14-1.30) nor in M1 (0.83, 0.79-0.88). These results suggest that the differences in STP between M2 and M1 are primarily presynaptic in origin and are not confounded by postsynaptic effects (**Figure S1**).

The differences in STP may indicate differences in release probability (*p*_N_) that in turn could result from differences in the coupling distance (CD). As a first test for potential differences in CD, we analyzed the variability of synaptic delays (Bullmann et al., 2024; Schwarze et al., 2026). The absolute values of the synaptic delays were similar in both brain areas (M2: 2.9 ms, 2.5-3.3 ms; M1: 2.7 ms, 2.3-3.4 ms; **Figure 1E**). However, the variability of the delays was significantly larger in M2 (SD_delay_, 0.22 ms, 0.14-0.34 ms) than in M1 (0.09 ms, 0.07-0.10 ms; **Figure 1F**). This indicates that at PN synapses in M2 transmitter release is less reliably coupled to the timing of the action potential than in M1, which could indicate a difference in the coupling configuration.

To test for different postsynaptic glutamate receptor contributions between PN synapses in M2 and M1, we analyzed the decay time constants (τ_decay_) of EPSCs. Under comparable voltage-clamp conditions, a larger τ_decay_ may indicate a greater involvement of N-methyl-D-aspartate receptors (NMDAR), since they exhibit slower opening and closing kinetics than AMPAR. We found significantly larger τ_decay_ in M2 (4.4 ms, 3.1-6.0 ms) compared to M1 (1.9 ms, 1.2-4.7 ms; **Figure 1G**). Furthermore, strong and delayed depolarizing currents in M2 were recorded especially when raising the extracellular Ca^2+^ ([Ca^2+^]_e_) to 3 mM. Inhibition of NMDAR with 50 µM 2-Amino-5-phosphonovaleriansäure (APV) reduced these currents in M2 accompanied by a significant decrease in EPSC charge (0.53, 0.36-0.78, n=7), while EPSC amplitudes remained unaffected (0.88, 0.77-1.13; **Figure S2**). In M1 APV affected neither the amplitude (1.0, 1.0-1.06, n=5) nor the charge of EPSCs (1.0, 0.92-1.02). These findings suggest a contribution of postsynaptic NMDAR to the later phase of the EPSC in M2 but not in M1.

In summary, we found significant differences in STP between synapses in M2 and M1, accompanied by significant differences in the temporal synchronization of release with the action potential. The differences may be explained by different CDs between the VGCCs and the release sensor, different *p*_N_ and/or differences in vesicle pool size and replenishment.

### Tight coupling between Ca^2+^ channels and release sensor in M2 and M1

To directly test for differences in CDs between M2 and M1 we analyzed the effects of ethylene glycol-bis(β-aminoethyl ether)-N,N,N′,N′-tetraacetic acid (EGTA) on release. In moderate concentrations, this kinetically slow Ca^2+^ chelator only interferes with release in loose coupling regimes but not if coupling is tight. It is thus used as a standard indicator to differentiate between tight and loose coupling (Naraghi and Neher, 1997; Eggermann et al., 2011; Bornschein and Schmidt, 2018). To compare the EGTA sensitivity of release at L2/3-L5PN synapses in M2 and M1 we examined EPSC amplitudes evoked by extracellular stimulation before, during and after incubation with a cell-permeant tetra(acetoxymethyl ester) of this Ca^2+^ chelator (EGTA-AM) (**Figure 2A**). Following 10 min of stable baseline recording, slices were perfused with ACSF containing either 10 µM EGTA-AM or the solvent DMSO/Pluronic alone. Incubation with the AM compound for 30 min allows accumulation of EGTA inside the cells (Atluri and Regehr, 1996; Matsui and Jahr, 2003; Hefft and Jonas, 2005) and was terminated by subsequent perfusion with standard ACSF for another 10 min (test period). EGTA effects were quantified during the test period and normalized to the corresponding baseline values. As expected, L2/3-L5PN synapses in M1 were insensitive to 10 µM EGTA-AM and showed no significant reduction in EPSC amplitudes (1.08, 0.92-1.17, n=6) compared to control recordings (0.97, 0.81-1.19, n=7; **Figure 2B**). Surprisingly, no EGTA sensitivity was detected in M2 synapses either (0.86, 0.79-0.90, n=7). This indicates that the larger SD_Delay_ in EPSC amplitudes in M2 is due to factors other than loose coupling. To test for proper functioning of the chelator-AM method, we also performed these experiments at L2/3-L5PN synapses in the PFC. Release from these synapses was previously shown to be sensitive to 10 µM EGTA-AM (Schwarze et al., 2026). We could confirm the significant amplitude reduction after application of 10 µM EGTA-AM here (0.46, 0.37-0.48, n=5), providing a positive control for the proper functioning of the EGTA-AM method.

**Figure 2.**
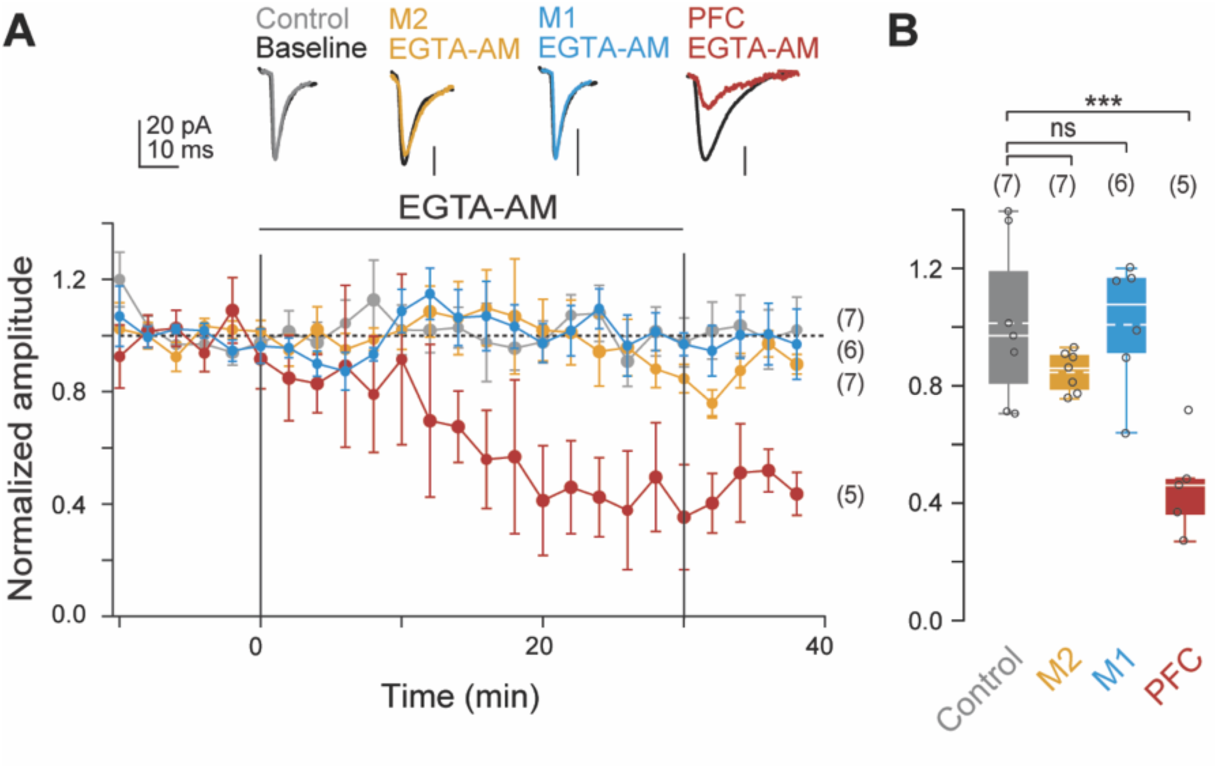
Low EGTA-AM sensitivity reveals tight coupling in M2 and M1. (**A**) Baseline-normalized, averaged EPSC amplitudes (means±SEMs, 2-min bins) recorded from layer 5 pyramidal neurons (L5PNs) after extracellular stimulation in layer 2/3 in different cortical regions (M1, M2, PFC). After stable baseline recordings for 10 min, 10 µM EGTA-AM were applied for 30 min in M1 (blue), M2 (orange), and PFC (red). Control recordings were performed in M2 by application of the solvent DMSO/Pluronic without EGTA (gray). Insets show averaged example EPSCs during baseline recordings (black, -10 to 0 min) and rinse with ACSF (test period, 30-40 min) for each group. Following EGTA-AM application, EPSC amplitudes remained stable in M1 and M2, compared to a marked decrease over time in PFC. (**B**) Box plots comparing the EPSC amplitudes after EGTA-AM application (test period) normalized to baseline values (dots represent individual experiments). EPSC amplitudes were not significantly affected by EGTA application in M1 (P=1.0, one-way ANOVA) and M2 (P=0.425) compared to control recordings suggesting tight coupling in contrast to loose coupling in PFC (***P<0.001).

These findings indicate that the CD in M1 synapses is tight, which is in line with the high temporal synchronization between presynaptic AP and release. Although transmitter release was temporally less precise in M2, these synapses were equally insensitive to EGTA, indicating that they also operate with tight Ca²⁺-influx-to-release coupling.

### High release probability in M2 and M1

To test whether different *p*_N_ can account for the observed differences in STP between L2/3-L5PN synapses in M2 and M1 we performed multiple probability fluctuation analysis (MPFA) (Clements and Silver, 2000). EPSC amplitudes were recorded at different extracellular Ca^2+^ concentrations ([Ca^2+^]_e_) and parabolas were fitted to the variance-mean plots of M2 and M1 synapses (**Figure 3A, B**). These fits yielded similarly high *p*_N_ values for synapses in M2 (0.69, 0.60-0.71, n=8) and M1 (0.68, 0.67-0.70, n=6; **Figure 3C**).

**Figure 3.**
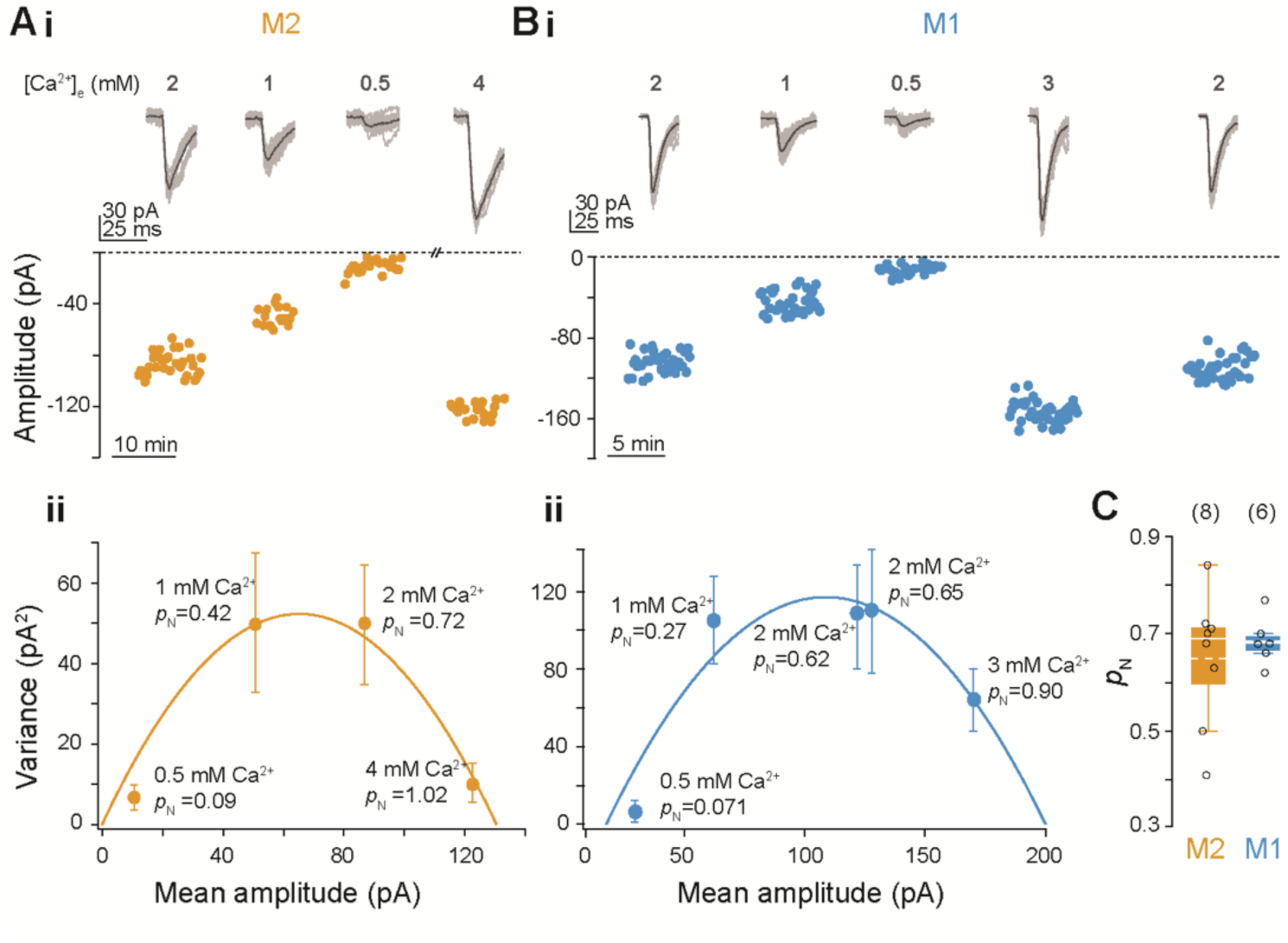
Release probability is high in M2 and M1. (**A, B**) Multiple-probability fluctuation analysis (MPFA) performed in synapses between L2/3 and L5PNs in M2 (**A**) and M1 (**B**), respectively. **i.** Example recording of EPSC amplitudes at different extracellular calcium concentrations ([Ca²⁺]_e_ in mM: 0.5, 1, 2, 3 and/or 4). Representative traces above show averaged EPSCs for each condition. **ii.** Variance versus mean amplitude plots (with error bars indicating the variance of the variance), where solid lines represent parabolic fits to estimate vesicular release probability (*p*_N_). Each point represents a different [Ca²⁺]_e_ condition. (**C**) Box plot comparing *p*_N_ at 2 mM [Ca²⁺]_e_ between M2 (orange) and M1 (blue). Note, that *p*_N_ is similarly high in M2 and M1 (P=0.547, MWU).

In summary, the differences in the PPRs which we observed between M2 and M1 cannot be explained by differences in *p*_N_. These finding suggest that other mechanisms, in particular the recruitment of vesicles to the ready-releasable pool (RRP), are responsible for the PPF at M2 synapses.

### Increased vesicle pool size and accelerated vesicle replenishment in M2

Recent studies show that an important determinant of STP is the kinetics of vesicle replenishment, which can even lead to a reversal overfilling of the RRP and to pronounced facilitation (Doussau et al., 2017; Weichard et al., 2023). Therefore, in the next step we investigated whether differences in vesicle pool size and reloading kinetics can account for the observed differences in STP. To analyze vesicle recruitment we induced a strong depression by sustained high-frequency (HF) synaptic activity, thereby, driving the synapses into a steady-state between release and replenishment, which is necessary for cumulative amplitude analyses (Schneggenburger et al., 1999; Neher, 2015). HF depression of EPSC amplitudes was induced by extracellularly stimulating L2/3-L5PN synapses with 50 pulses at 33 Hz in M2 and M1 (**Figure 4A, B**). Cumulative EPSC amplitudes were calculated and normalized to the first amplitude in the train (A_1_) to correct for the variability in the number of activated fibers and synapses between cells and runs during repeated fiber tract stimulations (**Figure 4C**). The slope of the linear fit to the late phase of the cumulative amplitude plot is a measure of the steady-state recruitment rate and the back-extrapolated y-intercept (y_0_) gives an approximation for the size of the vesicle pools contributing to release during the train (Schneggenburger et al., 1999). y_0_ not only includes the RRP in the narrower sense but also at least parts of the intermediate replenishment pool (RP; (Brachtendorf et al., 2025); see Methods for details). The slope of the cumulative amplitude plot was similar for M2 and M1 connections (M2: 0.87 pA/ms, 0.50-1.28 pA/ms; M1: 0.86 pA/ms, 0.58-1.12 pA/ms; **Figure 4D**), suggesting similar replenishment kinetics of the RP which is the rate-limiting step during steady-state vesicle recruitment. Furthermore, we found larger normalized y_0_ values in M2 (10.25, 8.36-12.88, n=14) compared to M1 connections (6.70, 5.37-6.97, n=10). In the presence of replenishment y_0_ underestimates the vesicle pool size and can be corrected according to Neher 2015 (M2: 14.90, 11.95-16.65; M1: 7.88, 6.89-9.18; see Methods and **Figure S3**) (Neher, 2015). To determine the release probability per vesicle, the first EPSC amplitude was divided by the size of RP and RRP (A_1_/y_0_) and designated as release fraction. This release fraction was smaller in M2 (0.10, 0.08-0.12; corrected 0.07, 0.06-0.09) compared to M1 (0.15, 0.14-0.19; corrected 0.13, 0.10-0.15). In summary, the size of RRP and RP was significantly larger during steady-state vesicle recruitment in M2 compared to M1; consequently, the release fraction was smaller in M2.

**Figure 4.**
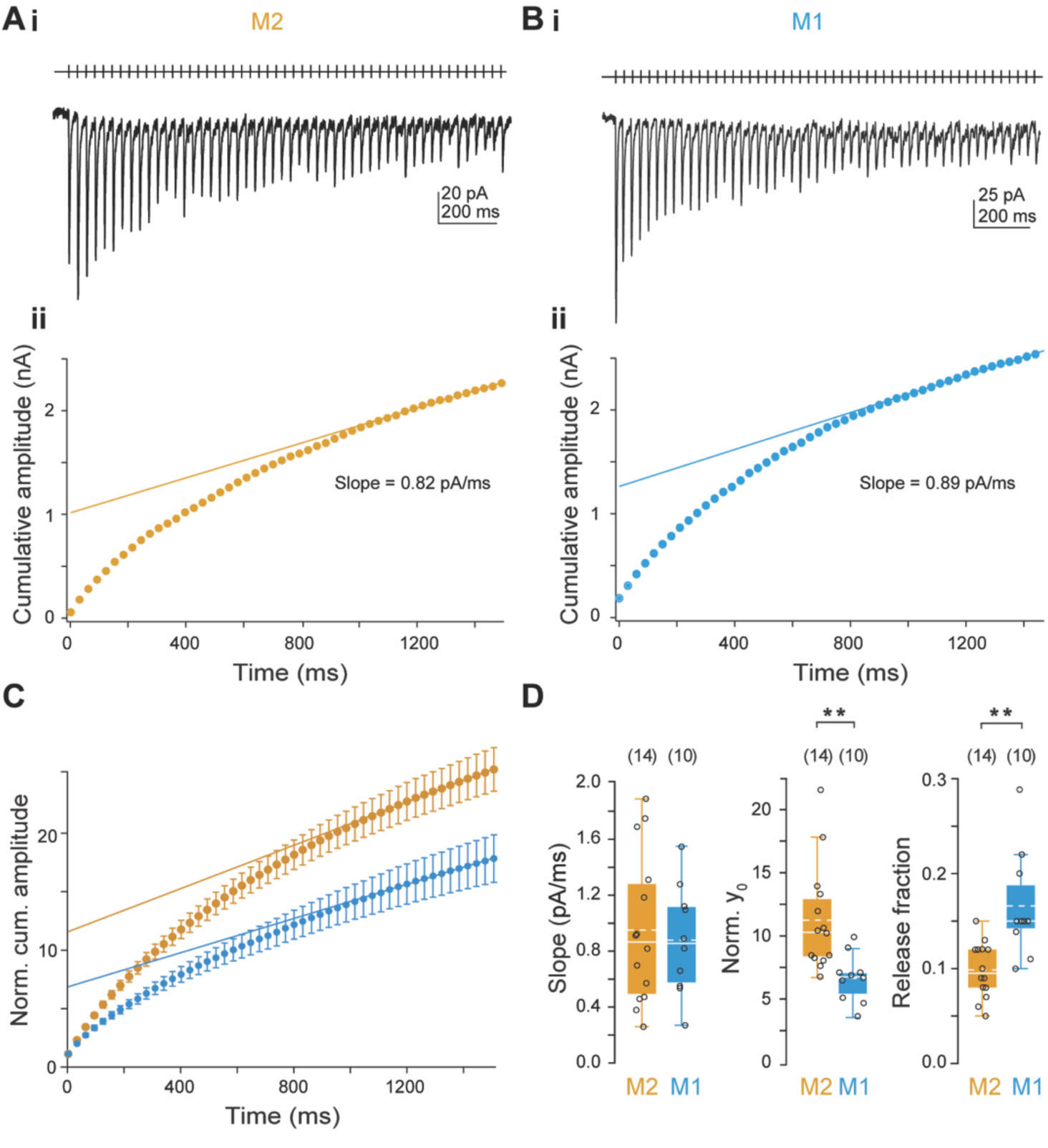
Cumulative postsynaptic responses to high-frequency stimulation in M2 and M1. (**A, B**) **i.** Examples of averaged EPSC trains (9 repetitions each) elicited via extracellular stimulation in L2/3 at 33 Hz and recorded in M2 (**A**) and M1 L5PNs (**B**). The high-frequency (HF) stimulation protocol, shown above the traces, was identical for both regions. **ii.** Cumulative amplitude plots calculated from the EPSCs in **i** with back-extrapolated linear fits to the steady-state phases (40^th^ to 50^th^ amplitude) to derive y-intercepts (y_0_) and slopes. (**C**) Averaged cumulative EPSC amplitudes (means ± SEMs) from M2 (orange) and M1 (blue) neurons with back-extrapolated linear fits to the steady-state phases normalized to the first amplitude in the train. (**D**) Summary of slopes (left), normalized y_0_ (middle), and release fractions (A_1_/y_0_, right) in M2 and M1. While the slope showed no significant difference between the two regions (P=0.861), M2 neurons exhibit significantly higher normalized y_0_ (**P=0.002, MWU) and lower release fractions (**P=0.003) compared to M1 neurons.

To further investigate the size of the contributing pools and the kinetics of vesicle recruitment we proceeded by analyzing the decrease of EPSC amplitudes during HF depression and the recovery from depression in M2 and M1 connections in more detail. The time course of EPSC amplitudes during HF stimulation (50 pulses at 33 Hz) was analyzed based on the recordings in **Figure 4** after normalization to the first amplitudes in the train (**Figure 5A, B**). We found that HF depression was best described by biexponential functions in both brain regions (for evaluation criteria see Methods and **Figure S4**). The biexponential decay of EPSC amplitudes supports the presence of an intermediate RP between the RRP and an infinite reserve pool (RSP) (Weichard et al., 2023; Brachtendorf et al., 2025). Assuming this pool arrangement, the relative contribution of exponential components provides information about the size of RRP and RP, and their time constants describe the kinetics of vesicle recruitment to RRP and RP. During initial HF depression τ_fast_ was significantly slower in M2 (0.07 s, 0.05-0.08 s, n=13) compared to M1 (0.03 s, 0.02-0.05 s, n=10; **Figure 5C**), whereas the exponential component A_1_ was similar in both brain regions (M2: 0.43, 0.21-0.47; M1: 0.46, 0.36-0.52) suggesting that the replenishment of the RRP (r_1_) is faster in M2 whereas the amount of vesicles released from the RRP (A_1_) is similar in M2 and M1. HF depression continues with similar τ_slow_ values in both brain regions (M2: 0.53 s, 0.44-0.96 s; M1: 0.54 s, 0.29-0.71 s), but with a significantly larger A_2_ component in M2 (0.56, 0.42-0.68) in comparison to M1 (0.39, 0.33-0.49), indicating a similar replenishment rate of the RP (r_2_) but the presence of a larger RP (A_2_) in M2. HF depression terminates at similar relative steady-state amplitudes (A_ss_) in M2 (0.21, 0.11-0.29) and M1 (0.20, 0.13-0.25).

**Figure 5.**
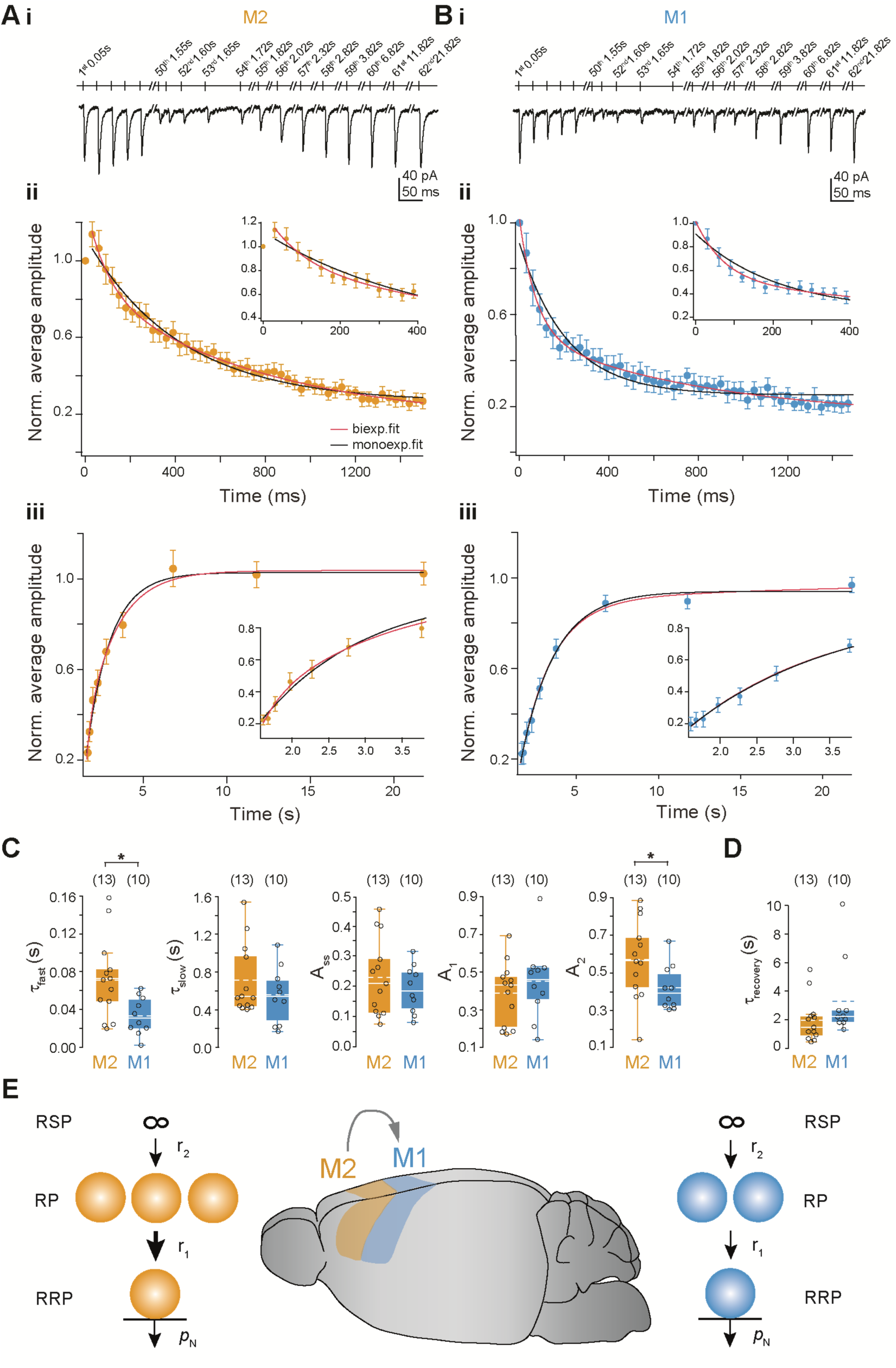
Vesicle replenishment is faster in M2 compared to M1 synapses. (**A, B**) **i.** Representative EPSC traces recorded from M2 (**A**) and M1 (**B**) L5PNs in response to extracellular stimulation in L2/3. Stimulation protocols (indicated above) with increasing interstimulus intervals (ISIs) after the HF train were used to record recovery from depression. **ii.** Normalized averaged EPSC amplitudes of the first 50 stimuli plotted over time. Data were fitted with monoexponential (black) and biexponential (red) functions to describe the kinetics of HF synaptic depression. Insets show the initial decay phase on an expanded timescale (0-400 ms). The decrease in EPSC amplitudes was clearly biexponential (M2: χ²_mono_/χ²_bi_=3.41; M1: χ²_mono_/χ²_bi_=4.91) and revealed significantly larger initial time constants (τ_fast_) in M2 compared to M1, indicating a slower vesicle depletion in M2. **iii.** Normalized averaged EPSC amplitudes of the recovery phase from steady state plotted as a function of recovery time and fitted with biexponential (red) and monoexponential (black) functions. Insets show magnified views of the initial recovery phase (1.65-3.8 s) which was predominantly biexpoential in M2 (χ²_mono_/χ²_bi_=1.72) and monoexponential in M1 (χ²_mono_/χ²_bi_ =1.00; cf. Figure S4). (**C**) Box plots summarizing kinetic parameters derived from biexponetially fitted HF depression: fast and slow decay time constants (τ_fast_, τ_slow_), steady-state amplitude (A_ss_), and amplitude coefficients (A_1_, A_2_). Recordings from M2 neurons show significantly larger τ_fast_ (*P=0.020, MWU) and A_2_ values (*P=0.032) compared to M1, whereas other parameters do not differ significantly between the two regions (P=0.598, P=0.642, P=0.515). (**D**) Box plot showing recovery time constants (τ_recovery_) expressed as weighted τ derived from mono- and biexponential fits to the recovery from depression in the M1 and M2 region. τ_recovery_ is not significantly faster in M2 than in M1 synapses (P=0.088, MWU). (**E**) Schematic representation of the vesicle pools in M2 (left, orange) and M1 (right, blue) as well as a mouse brain diagram showing the corresponding anatomical localization of the M2 and M1 regions within the cerebral cortex (middle). In the sequential pool model synaptic vesicles are released from the readily releasable pool (RRP) with *p*_N_. Replenishment occurs from an intermediate replenishment pool (RP) and an infinite reserve pool (RSP) with rate constants r_1_ and r_2_. Our findings suggest that in M2 the RP is larger and replenishment of the RRP is much faster than in M1 corresponding to the significant larger τ_fast_ and A_2_ in (**C**).

Additionally, we examined the recovery from depression by application of extracellular stimuli with increasing intervals at the end of the HF train. The amplitude recovery was fitted either with a mono- or a biexponential function with either one or two time constants. Recovery was biexponential in 8 out of 13 recordings in M2 and in 4 out of 10 recordings in M1 (**Figure S4**). For comparison of recovery kinetics a weighted time constant (τ_recovery_) was calculated from τ_fast_ and τ_slow_ of biexponential fits (see Methods). τ_recovery_ was not significantly different in M2 (1.46, 0.92-2.22) and in M1 (2.19, 1.83-2.63; **Figure 5D**).

These results indicate faster vesicle recruitment from a larger RP in M2 connections, which could explain PPF (**Figure 5E**). In contrast, paired-pulse depression in M1 synapses could be caused by a depletion of the vesicle pool.

## Discussion

Our data show PPF at glutamatergic synapses from L2/3 to L5PNs in M2 as opposed to PPD in M1 (**Figure 1**). Despite this difference in STP we found no evidence of differences in the EGTA sensitivity of release or in *p*_N_ between areas (**Figure 2, 3**). Our results suggest that the differences in STP arise from a larger RP in M2 synapses in comparison to M1, while the size of the RRP was similar (**Figure 4, 5**).

The insensitivity of release to low to moderate concentrations of EGTA is a standard indicator of tight Ca^2+^-influx-to-release coupling (Adler et al., 1991; Naraghi and Neher, 1997; Bucurenciu et al., 2008; Eggermann et al., 2011; Bornschein and Schmidt, 2018; Bornschein et al., 2019). Thereby, *p*_N_ not solely depends on the CD between VGCCs and the sensor, but also on the size of the Ca^2+^ influx as well as the kinetics and sensitivity of the release machinery. We assume the same Ca^2+^ sensor affinity in M2 and M1, since PN synapses in neocortex including both motor areas express Syt1 as the predominating Ca^2+^ sensor of release (Berton et al., 1997; Zhang et al., 2014; Bornschein et al., 2025). Differences in EGTA sensitivity are unlikely to result from differences in Ca^2+^ sensor affinity. Previous studies indicated that even if the sensor’s k_on_ value and its affinity should differ 4 to 10 fold, the CD would have the major impact on EGTA sensitivity (Schneggenburger and Neher, 2000; Bollmann et al., 2000; Kusch et al., 2018; Bornschein et al., 2019; Bornschein et al., 2025). In addition, we assume that Ca^2+^ influx is similar. This assumption appears justified since in previous studies no differences were observed between PFC and S1 and also not between young and mature synapses in S1 (Baur et al., 2015; Bornschein et al., 2019; Schwarze et al., 2026). Together, these observations suggest that PN synapses in both M1 and M2 operate with similarly tight coupling and comparably high initial *p*_N_. Besides similar EGTA sensitivity, we found a significantly larger SD_Delay_ at M2 than at M1 synapses, indicating that factors other than coupling distance contribute to the temporal precision of transmitter release. Potential determinants include the organization and gating of release-triggering VGCCs (Meinrenken et al., 2002; Fedchyshyn and Wang, 2005), the presynaptic AP waveform (Sabatini and Regehr, 1997; Borst and Sakmann, 1999; Chao and Yang, 2019), endogenous Ca^2+^ buffers (Eggermann and Jonas, 2011) or the heterogeneity in vesicle priming states (Lin et al., 2022). However, similar EGTA sensitivity and *p*_N_ argue against major differences in coupling architecture, Ca^2+^ influx, or release efficacy, while endogenous buffers are unlikely to play a major role because L5PNs express only low levels of mobile Ca^2+^ buffers (Helmchen et al., 1996; van Tran and Stricker, 2018). Nevertheless, the increased release jitter reflected by a larger SD_delay_ and a slower τ_decay_, which may only partly be due to a larger NMDA receptor contribution, suggest that M2 synapses are less specialized for the initial synchronous release process than M1 synapses.

We found significant differences in STP with short-term facilitation in M2 as opposed by short-term depression in M1. PPD was found previously in fast and highly reliable cortical PN synapses in S1 (Frick et al., 2008; Bornschein et al., 2020) emphasizing the first transmission process. Whereas, PPF was described at highly plastic synapses in PFC (Shin et al., 2025; Schwarze et al., 2026) and hippocampus (Vyleta and Jonas, 2014), emphasizing the subsequent release and providing a higher plasticity and flexibility for information processing. These findings indicate that coupling architecture and *p*_N_ alone are insufficient to predict the direction and magnitude of STP. Various other factors besides CD and *p*_N_ can influence STP, that are the number of occupied release sites (*N*_occ_), the replenishment of *N*_occ_ or the recruitment of newly formed *N*_occ_ as well as the expression of endogenous Ca^2+^ buffers (Neher, 1998; Felmy et al., 2003; Blatow et al., 2003; Matveev et al., 2004; Regehr, 2012) or facilitation sensors (Shin et al., 2025). Postsynaptic sources are unlikely to account for differences in STP, since the influence of receptor saturation and desensitization - as demonstrated by application of Kyn - is negligible (**Figure S1**) (Neher & Sakaba, 2001).

Our data suggest that the differences in STP result from differences in the size of the RP, from which vesicle recruitment occurs thereby affecting the speed and reliability of vesicular release, rather than from differences in *p*_N_. HF train analyses are commonly used to determine pool sizes and the kinetics of vesicle replenishment (Brachtendorf et al., 2025), although they cannot distinguish between parallel, sequential and the more recently proposed LS-TS pool models (Neher and Brose, 2018; Kusick et al., 2022; Neher, 2024; Brachtendorf et al., 2025). The biexponential decay of HF depression is consistent with either a series-connected finite replenishment pool (RP) or a subdivision of the RRP into parallel pools (Sakaba, 2006; Wölfel et al., 2007; Hallermann et al., 2010; Ritzau-Jost et al., 2018). Based on our previous work indicating an RP intercalated between the RRP and RSP at cortical PN synapses (Bornschein et al., 2019; Schwarze et al., 2026), which matures and increases during postnatal development and thereby increases PPR (Bornschein et al., 2020), we assume a sequential pool model at the same synapses in M2 and M1. In this model, STP is largely determined by the transition rate of synaptic vesicles from the RP to the RRP (Miki et al., 2016; Doussau et al., 2017; Schmidt, 2019), whereas sustained recruitment depends on the size of the RP. Accordingly, rapid replenishment can progressively overfill the initial RRP and increase N_occ_ above baseline, thereby promoting PPF (Valera et al., 2012; Brachtendorf et al., 2015; Miki et al., 2016; Doussau et al., 2017; Shin et al., 2025). In cumulative analysis, the larger y-intercept observed in M2 could reflect an enlarged RRP and/or RP, as it represents the sum of both pools in the sequential model and also includes unoccupied release sites (Brachtendorf et al., 2025). Together with the faster refilling kinetics, this suggests a highly dynamic vesicle cycle that favors rapid recruitment, potentially at the expense of maintaining a uniformly mature population of release-ready vesicles.

In summary, M1 exhibited properties similar to those of S1, indicating that PN synapses in primary cortices irrespective of sensory or motoric operate with tight nanodomain coupling and high *p*_N_ and emphasize the first release process. M2 had intermediate properties, on the one hand showing tight coupling and high *p*_N_ similar to those in the primary cortices and on the other hand facilitation reminiscent of higher order cortices, thereby supporting subsequent release and enhanced synaptic plasticity. These findings imply that PNs in intermediate areas operate with intermediate synaptic characteristics.

## Methods

### Slice Preparation

Patch-clamp recordings were performed on layer 5 pyramidal neurons (L5PNs) in acute brain slices taken from C57BL/6 mice of both sexes at postnatal days P21 to P24 (mature). The mice were decapitated under deep isoflurane (Curamed) inhalation anesthesia; the cerebrum was swiftly removed and immersed in cooled (0–4°C) artificial cerebrospinal fluid (ACSF), which contained the following concentrations (in mM): 125 NaCl, 2.5 KCl, 1.25 NaH_2_PO_4_, 25.6 NaHCO_3_, 1 MgCl_2_, 2 CaCl_2_, and 20 glucose, equilibrated with 95% O_2_ and 5% CO_2_ (pH 7.3–7.4). Unless otherwise specified, all chemicals were from SIGMA-Aldrich. Coronar acute brain slices (250 μm thick) were cut from M2 and M1 region (**Figure 1A**) using an HM 650 V vibratome (Microm) and maintained at 35°C - near physiological temperature - for 30 minutes. Subsequently, the slices were stored at room temperature (∼22°C).

Animal procedures were conducted in accordance with institutional guidelines for animal experiments, and were approved by the state directorate of Saxony, Germany.

### Electrophysiology

For experiments, cortical slices containing M1 or M2 regions were placed into a recording chamber, which was continuously perfused with 4 ml of artificial cerebrospinal fluid (ACSF) per minute, supplemented with 10 μM (−)-bicuculline methiodide (Tocris) and in some experiments with 50 µM 2-Amino-5-phosphonovaleriansäure (APV) and 0.25 mM kynurenic acid (Kyn) at a temperature of 33 ± 2°C. Patch pipettes, pulled from borosilicate glass (Hilgenberg) with a PC-10 puller (Narishige), exhibited final series resistances (R_s_) of approximately 5 MΩ for whole-cell recordings in acute brain slices. The patch pipettes were filled with the following pipette solution (in mM): 150 K-gluconate, 4 NaCl, 3 MgCl_2_, 3 Na_2_ATP, 0.3 NaGTP, 0.05 EGTA, 10 KHEPES, dissolved in purified water and adjusted to pH 7.3 with KOH.

Postsynaptic whole-cell patch-clamp recordings were performed under optical control (Zeiss Axioskop 2 FS plus) with an EPC9 amplifier and PatchMaster software (v2x92, HEKA). Excitatory postsynaptic currents (EPSCs) were recorded from visually identified L5PNs at a holding potential (V_m_) of −80 mV, corrected for the liquid junction potential of 16 mV. Data were filtered at 5 kHz and sampled at 10 kHz. During the recording, the series resistance (R_s_) and holding current (I_hold_) were constantly tracked. R_s_ compensation was dynamically adjusted every minute to achieve a remaining uncompensated R_s_ of 10-20 MΩ. Data were discarded if uncompensated R_s_ exceeded 30 MΩ or if I_hold_ dropped below −300 pA. Extracellular stimulation was applied in neocortical layer 2/3 using an ACSF-filled stimulation pipette to evoke EPSCs in L5PNs. A 200 µs bipolar pulse was administered, with stimulus intensities adjusted to achieve initial evoked EPSC amplitudes of approximately 100 pA, utilizing an ISO-Stim 01M (NPI Electronics) device. Extracellular stimulations were repeated every 10 seconds for up to 60 minutes.

To analyze chelator-AM effects on EPSC amplitudes (**Figure 2**) amplitudes were binned within 2 min intervals to calculate mean amplitudes (Baur et al., 2015). We used the well-established technique of utilizing the calcium chelator EGTA to interfere with neurotransmitter release (Naraghi and Neher, 1997; Nägerl et al., 2000). Exogenous calcium chelators were introduced into L2/3-L5PN synapses by applying their membrane-permeable AM forms to the bath solution (Atluri and Regehr, 1996; Matsui and Jahr, 2003; Hefft and Jonas, 2005). Following 10 min baseline recording in normal ACSF, slices were incubated for 30 min with either a control solution (0.1% DMSO and 0.01% Pluronic) or EGTA-AM in DMSO/Pluronic and afterwards reperfused with normal ACSF for a 10 min test period. The extended incubation time of 30 min allowed for substantial intracellular accumulation of the chelators (Tsien, 1981; Atluri and Regehr, 1996). This duration was carefully chosen to ensure sufficient chelator uptake while avoiding over-accumulation. The subsequent reperfusion with normal ACSF was crucial to remove excess extracellular chelators and establish a stable intracellular chelator concentration for the experiments.

Multiple-probability fluctuation analysis (MPFA) was performed to investigate *p*_N_ in L2/3-L5PN synapses (**Figure 3**). EPSC amplitudes were recorded at different extracellular Ca^2+^ concentrations ([Ca^2+^]_e_). To prevent postsynaptic receptor desensitization and saturation 0.25 mM Kyn were added to the bath solution (**Figure S1**). At [Ca^2+^]_e_ ≥ 3 mM we found strong NMDA-mediated post-depolarizations that impeded precise amplitude analyses (**Figure S2A**). Therefore, bath solutions were additionally supplemented with 50 µM of the selective antagonist APV that affected EPSC charges but not the amplitudes (**Figure S2B, C**). The MPFA approach applies binominal statistics and accounts for non-uniform quantal size (Quastel, 1997; Silver et al., 1998; Clements and Silver, 2000; Silver, 2003; Saviane and Silver, 2006). EPSC variances (σ²) were calculated according to Scheuss et al. (Scheuss et al., 2002) and plotted against mean EPSC amplitudes (I) and fitted with a parabolic function:

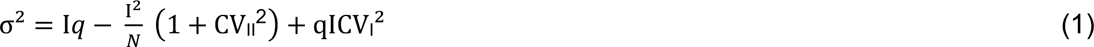

with *q* denoting quantal size, *N* representing the number of release sites or releasable vesicles, and CVI and CVII being coefficients of intrasite and intersite quantal variability, set at 0.3 (Clements and Silver, 2000). The variance of the variance was calculated according to (Meyer et al., 2001).

To investigate vesicle pool depletion and replenishment we evoked high-frequency depression by applying a repetitive stimulation pattern consisting of 50 stimuli delivered at 33 Hz, followed by 9 pulses at increasing intervals to examine recovery from depression *(70 ms, 100 ms, 200 ms, 300 ms, 500 ms, 1 s, 3 s, 5 s, 10 s;* **Figure 4, 5**). Through sustained high-frequency (HF) synaptic activity, the synapses reach a steady-state between release and replenishment, from which they slowly recover. During the 33 Hz train a cumulative analysis of EPSC amplitudes was performed. The slope of a line fit to the late phase of the cumulative amplitude (last 10-20 stimuli) provides an estimate of the steady-state recruitment rate, and the back-extrapolated y-intercept (y_0_) reflects a quantity close to the decrement of the vesicle pool, from which vesicles are released during the train (Schneggenburger et al., 1999; Neher, 2015). Assuming that during synaptic activity at mature L5PN synapses the ready-releasable pool (RRP) is replenished by SVs from a finite-size replenishment pool (RP), that is intermediate between the RRP and the infinite reserve pool (RSP) (Miki et al., 2016; Doussau et al., 2017; Schmidt, 2019), y_0_ represents the decrement of the RP and the RRP (Bornschein et al., 2020; Brachtendorf et al., 2025). In the presence of replenishment y_0_ underestimates the decrement in vesicle pool size and can be corrected using the following formula (**Figure S3**) (Neher, 2015):

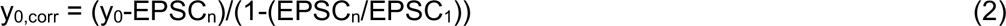

with EPSC_1_ and EPSC_n_ being the first and the last (n) EPSC amplitude in the train, respectively. The time course of HF depression was analyzed during the 33 Hz train and recovery from depression after the end of the train between 1.65 and 3.8 s (53^rd^ to 62^nd^ pulse). The protocol was repeated 10 times with a 10 s interval in between. EPSC amplitudes during HF depression and recovery phases were averaged, normalized to the first EPSC in the train and fitted with mono- and biexponential functions, respectively. The fit was accepted as bi-exponential if three criteria were fulfilled: (1) The sum of the squared differences between the experimental trace and the fit was at least 4% larger with the mono- compared to the biexponential fit (χ^2^_mono_/χ^2^_bi_>1.04). (2) The relative contribution of the fast and the slow component was >10% (e.g. 0.1<A_1_<0.9). (3) The fast and slow time constants differed by a factor >3. If one of the criteria was not met, a monoexponential fit was used instead (**Figure S4**) (Eshra et al., 2021). A biexponential decay of EPSC amplitudes during HF activity supports the assumption of a sequential pool model with an intermediate RP between RRP and RSP, whereas a monoexponential decay would occur when the RRP is recruited directly from the RSP (Brachtendorf et al., 2025). Analyses of biexponential components of HF depression reveals the relative contribution of RRP (A_1_) and RP (A_2_), while the time constants τ_fast_ and τ_slow_ describe the kinetics of vesicle recruitment to RRP (r_1_) and RP (r_2_), respectively (Brachtendorf et al., 2025).

To compare mono- and biexponential recovery kinetics a weighted time constant (τ_recovery_) was calculated for biexponential fits (**Figure 5D**) (Weichard et al., 2023):

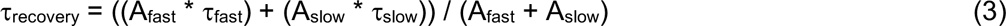

with τ_fast_ and τ_slow_ being the fast and slow time constants, and A_fast_ and A_slow_ are the amplitudes of the two exponential components, respectively.

### Quantification and statistical analysis

Electrophysiological data were analyzed using custom-written routines in Igor Pro 8 (version 8.04, WaveMetrics). Data are presented as median and interquartile range (IQR), mean ± standard deviation (SD), and mean ± standard error of the mean (SEM), respectively. Within box plots boxes indicate medians and IQRs, dashed lines represent means, whiskers extend to the 10^th^ and 90^th^ percentiles and data points falling outside this range are depicted as individual outliers. The number of experiments (n) is indicated in parentheses.

Wilcoxon signed-rank test (WSR) was used to compare pre- and post-treatment data. Mann-Whitney rank-sum test (MWU) was applied for unpaired comparisons. For multiple comparisons a One-Way ANOVA was performed after ensuring normal distribution using Shapiro–Wilk test. A post-hoc test (Tukey’s Honest Significant Difference test, HSD) was used to identify specific group differences.

All statistical analyses were performed using Python version 3.11.7 along with the following libraries: NumPy (1.26.4) for numerical computations and array handling, Pandas (2.1.4) for data handling and preprocessing, Matplotlib (3.8.0) and Seaborn (0.12.2) for data visualization, SciPy (1.11.4) for statistical testing. All statistical tests were two-tailed, with significance levels denoted as follows *P≤0.05, **P≤0.01, ***P≤0.001, and ****P≤0.0001. The number of experiments (n) represents individual cells and is chosen high enough to ensure result consistency and facilitate appropriate statistical inference.

## Acknowledgements

We thank Gudrun Bethge for technical assistance.

This work was supported by a grant of the German research foundation (DFG SCHM1838/6-1 to HS).

## Additional information

### Author contributions

Conceptualization: G.B., H.S.; Funding acquisition: H.S.; Methodology: all authors; Investigation: A.Y., G.B., H.S.; Data Analysis: A.Y., G.B., S.B., H.S.; Visualization: A.Y., G.B., S.B.; Resources: H.S.; Supervision: G.B., H.S.; Writing – original draft: A.Y., G.B.; Writing – review & editing: all authors.

### Data and code availability

All data are available in the main text or in the supplemental information and will be shared by the corresponding author upon request.

## Additional files

### Supplemental Material

Document S1. Figures S1–S4

