## Supplemental Material S1 for "Pyramidal neuron synapses in M2 exhibit properties intermediate between prefrontal cortex and M1 synapses"

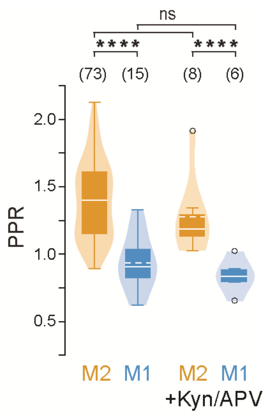


**Figure S1. PPR is not influenced by postsynaptic effects.**

Summary of PPRs recorded from L2/3-L5PNs at 50 Hz in M2 and M1. Postsynaptic saturation and desensitization were inhibited by 0.25 mM Kyn and NMDARs were blocked by 50 µM APV. PPR was not significantly altered in M2 (P=0.220, MWU) and M1 (P=0.470) compared to untreated recordings indicating no influence of postsynaptic effects on STP.

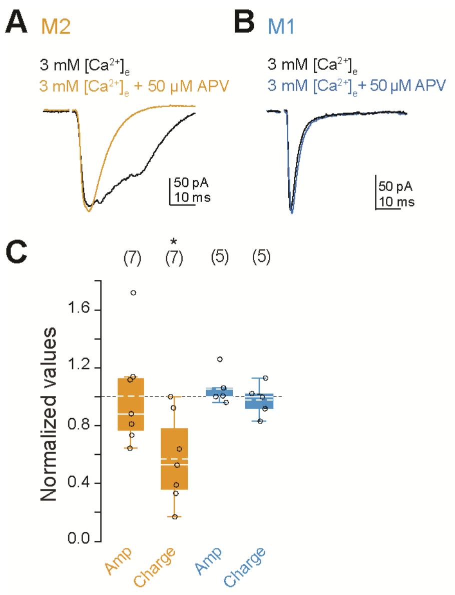


**Figure S2. Determination of a NMDA-mediated EPSC component in M2 rather than in M1 neurons.**

(**A, B**) Example recordings from L2/3-L5PN connections in M2 (**A**) and M1 (**B**), respectively, in 3 mM extracellular $Ca^{2+}$ ($[Ca^{2+}]_e$). To block postsynaptic NMDA receptors in M2 (orange) and M1 (blue) 50 µM APV were added to the extracellular solution.

(**C**) Quantification of EPSC amplitudes (Amp) and charges in M2 (orange) and M1 (blue), normalized to their respective baseline values (3 mM $[Ca^{2+}]_e$, 0 mM APV). APV significantly reduced postsynaptic charges in M2, rather than EPSC amplitudes (P=0.688, *P=0.028, P=0.285, P=0.854, WSR).

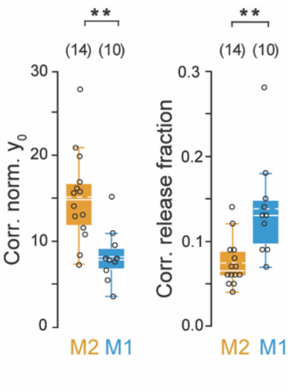


**Figure S3. Correction of vesicle pool size for replenishment.**

Summary of corrected normalized y-intercepts ($y_0$, *left*) and corrected release fractions ($A_1/y_0$, *right*) in M2 and M1. M2 neurons exhibit significantly higher normalized $y_0$ (**P=0.002, MWU) and lower release fractions (**P=0.002) compared to M1 neurons.

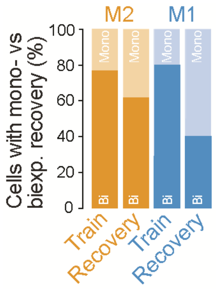


**Figure S4. Mono- vs. biexponential HF depression and recovery of EPSC amplitudes in M2 and M1.**

Proportion of cells exhibiting a mono- or biexponential decrease in EPSC amplitudes during high-frequency (HF) stimulation and a mono- or biexponential increase during the subsequent recovery from synaptic depression. The time course of HF depression is predominantly biexponential in both M2 and M1. During the recovery phase, the majority of M2 connections display a biexponential recovery from depression, whereas EPSC amplitudes in M1 increase mainly monoexponential.
